# Genome-wide decoupling of H2Aub and H3K27me3 in early mouse development

**DOI:** 10.1101/2021.03.23.436550

**Authors:** Yezhang Zhu, Jiali Yu, Yan Rong, Yun-Wen Wu, Heng-Yu Fan, Li Shen

## Abstract

Polycomb group (PcG) proteins are crucial chromatin regulators during development. H2Aub and H3K27me3 are catalyzed by Polycomb-repressive Complex 1 and 2 (PRC1/2) respectively, and largely overlap in the genome due to mutual recruitment of the two complexes. However, whether PRC1/H2Aub and PRC2/H3K27me3 can function independently remains obscure. Here we uncovered a genome-wide decoupling of H2Aub and H3K27me3 in preimplantation mouse embryos, at both canonical PcG targets and broad distal domains. H2Aub represses future bivalent genes without H3K27me3 but does not contribute to maintenance of H3K27me3-dependent non-canonical imprinting. Our study thus revealed their distinct and independent functions in early mammalian development.

## Introduction

Mono-ubiquitination of histone H2A at lysine 119 (H2AK119ub1 or H2Aub) and trimethylation of histone H3 at lysine 27 (H3K27me3) are deposited by Polycomb-repressive Complex 1 (PRC1) and PRC2, respectively. The two intimately associated histone modifications play central roles in Polycomb group protein (PcG)-mediated transcriptional repression and are of paramount importance to mammalian development (Faust et al. 1998; O’Carroll et al. 2001; Pasini et al. 2004; Posfai et al. 2012; Schuettengruber et al. 2017). While the classic model suggested recruitment of PRC1 by PRC2-mediated H3K27me3 deposition (Cao et al. 2002; Wang et al. 2004), recent studies also demonstrated the recruitment of PRC2 by PRC1-mediated H2Aub deposition (Blackledge et al. 2014; Blackledge et al. 2020; Tamburri et al. 2020), suggesting a reciprocal recognition of H2Aub and H3K27me3 by PRC2 and PRC1 respectively. Indeed, PRC1/H2Aub and PRC2/H3K27me3 have been reported to largely overlap in the genome, particularly at canonical PcG targets (i.e., promoters of bivalent genes) (Boyer et al. 2006; Ku et al. 2008; Blackledge et al. 2015; Blackledge et al. 2020; Tamburri et al. 2020; Zepeda-Martinez et al. 2020). However, whether they repress PcG target genes cooperatively or redundantly is still under debate (Blackledge et al. 2015; Schuettengruber et al. 2017; Cohen et al. 2021), and it is largely obscure if they may also function independently at different genomic regions in some biological context.

In addition to decorating canonical PcG targets, H3K27me3 has also been reported to form broad distal domains in oocytes (Liu et al. 2016; Zheng et al. 2016), which are inherited by early embryos to mediate DNA methylation-independent non-canonical imprinting (Inoue et al. 2017; Matoba et al. 2018; Zhang et al. 2019). Remarkably, ectopic removal of H3K27me3 resulted in loss of non-canonical imprinting, suggesting that H3K27me3 plays a dominant role in silencing the maternal allele of non-canonical imprinted genes (Inoue et al. 2017). However, due to the interdependent recruitment of PRC1 and PRC2, it is still elusive whether H2Aub also contributes to this DNA methylation-independent non-canonical imprinting system.

Here we developed an ultra-sensitive ChIP-Seq method and generated allelic H2Aub profiles in mouse gametes and early embryos. Comparative analysis showed that oocytes exhibit similar distributions of H2Aub and H3K27me3 at both canonical PcG targets and broad distal domains. However, in early embryos, H2Aub instead of H3K27me3 is enriched at PcG targets, and H3K27me3 is only associated with broad distal domains. These observations indicate that H2Aub represses future bivalent genes without H3K27me3 but does not contribute to maintenance of H3K27me3-dependent non-canonical imprinting. Our results thus not only revealed an unexpected decoupling of H2Aub and H3K27me3 in early embryos, but also suggested their distinct and independent roles during preimplantation development.

## Results and Discussions

### H2Aub exhibits a similar landscape to H3K27me3 with broad distal domains in mouse oocytes

To profile H2Aub in mouse oocytes and early embryos, we first adopted the ultra-low-input micrococcal nuclease-based native ChIP-Seq (ULI-NChIP-Seq) protocol, which has been successfully used to map H3K4me3 and H3K27me3 from hundreds of cells including early embryos (Brind’Amour et al. 2015; Liu et al. 2016). However, when used with the H2Aub antibody, ULI-NChIP-Seq failed to generate a robust result from 500 mouse embryonic stem cells (mESCs) as it did for H3K4me3. We thus developed an improved ChIP-seq method, termed CATCH-Seq (<u>c</u>arrier DNA <u>a</u>ssisted <u>Ch</u>IP-Seq), by introducing to ULI-NChIP-Seq a synthetic double-stranded DNA carrier that can be completely removed during library amplification to reduce DNA loss (Fig. S1A, S1B). Remarkably, CATCH-Seq could produce high-quality H3K4me3 maps from as few as 20 mESCs (Fig.S1C). We also successfully mapped H2Aub with CATCH-Seq from 500 mESCs and obtained comparable results to published data of bulk cells (Fig. 1A).

**Figure 1.**
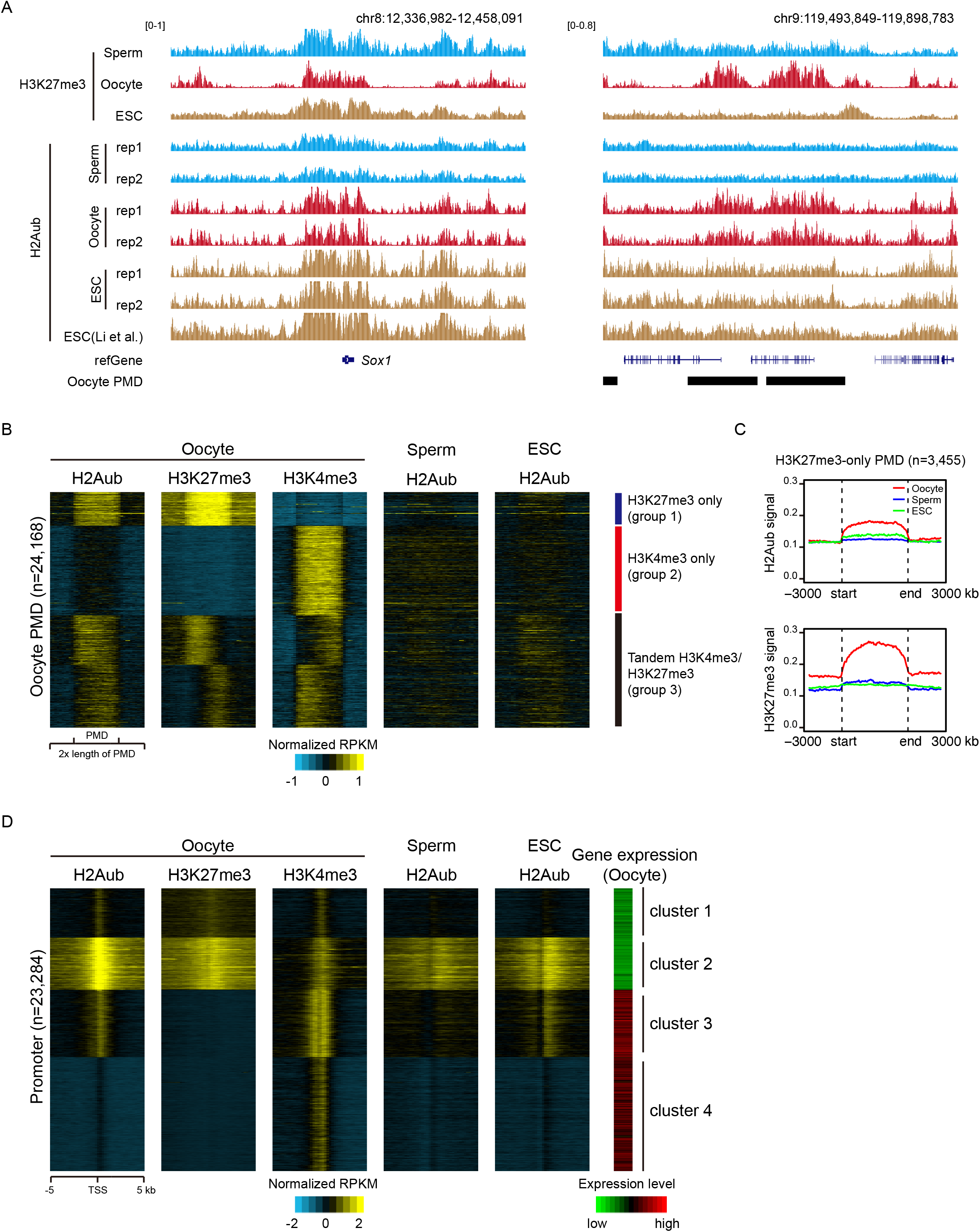
H2Aub exhibits a similar landscape to H3K27me3 with broad distal domains in mouse oocytes. **(A)** Genome browser view showing H2Aub and H3K27me3 enrichment in oocyte, sperm, and mESC near *Sox1* (left) and oocyte PMDs (right). **(B)** Heatmaps showing the enrichment of H2Aub, H3K27me3, and H3K4me3 around oocyte PMDs in oocyte, sperm, and mESC for various classes of PMDs. **(C)** Average H2Aub and H3K27me3 signals across H3K27me3-only PMDs in oocyte, sperm, and mESC. **(D)** Heatmaps showing the enrichment of H2Aub, H3K27me3, and H3K4me3 around transcription start sites (TSS) in oocyte, sperm, and mESC. Gene expression levels of corresponding genes in oocyte is also shown. Promoters are clustered into four groups using k-means clustering based on H2Aub and H3K27me3 signals in oocyte.

Next, we generated genome-wide landscapes of H2Aub in mouse oocytes and sperms with CATCH-Seq (Fig. 1A). Consistent with the largely overlapped distributions of H2Aub and H3K27me3 in mESCs (Blackledge et al. 2020; Tamburri et al. 2020; Zepeda-Martinez et al. 2020), H2Aub also exhibited similar genomic distribution to H3K27me3 in oocytes (Fig. 1A). As exemplified by genome browser views, H2Aub and H3K27me3 co-occupied not only canonical PcG targets (i.e., promoters of bivalent genes), but also the non-canonical broad distal H3K27me3 domains that are unique for oocytes and early embryos (Liu et al. 2016; Zheng et al. 2016) (Fig. 1A). In mouse oocytes, the broad H3K27me3 domains preferentially overlap with poorly methylated non-transcribing regions termed partially methylated domains (PMDs) (Zheng et al. 2016), we thus examined H2Aub enrichment in three previously identified subgroups of oocyte PMDs (Fig. 1B). This analysis revealed that the deposition of H2Aub within different PMDs highly resembled that of H3K27me3 in oocytes, with enrichment in H3K27me3 marked PMDs or H3K27me3/H3K4me3 co-marked PMDs (group 1 and 3) and depletion from H3K4me3 marked PMDs (group 2), which were not observed in mESCs and sperms (Fig. 1B, 1C). Notably, while H3K27me3 resides in H3K27me3/H3K4me3 co-marked PMDs in a non-overlapping manner (i.e., tandem distribution of H3K27me3 and H3K4me3) (Zheng et al. 2016) (Fig. 1B), H2Aub was less exclusive from H3K4me3 within these PMDs (Fig. 1B), suggesting that H2Aub was not directly involved in shaping the tandem but mutual exclusive distribution of H3K4me3 and H3K27me3 within these PMDs. We then further compared H2Aub, H3K27me3, and H3K4me3 enrichment at gene promoters by clustering all promoters into four groups based on their H2Aub and H3K27me3 signals in oocytes (Fig. 1D, S2A). Cluster 1 promoters showed weak enrichment of H2Aub, H3K27me3, and H3K4me3; Cluster 2 promoters exhibited strong association with H2Aub, H3K27me3, and H3K4me3, representing bivalent promoters (Fig. 1D, S2A, S2B). In contrast to transcriptionally silenced Cluster 1 and Cluster 2 promoters, Cluster 3 and Cluster 4 promoters showed active transcription and were associated with H3K4me3 but not H3K27me3 (Fig. 1D, S2A, S2C). Notably, Cluster 3 promoters, which exhibited the highest level of H3K4me3 enrichment, were also slightly enriched for H2Aub (Fig. 1D, S2A), in agreement with the reported association of non-canonical PRC1 with transcription activation (Aranda et al. 2015; Cohen et al. 2020). Consistently, while Cluster 2 promoters are bound by both CBX7 and RYBP, which represent canonical and non-canonical PRC1 respectively, in mESCs, Cluster 3 promoters are preferentially bound by RYBP (Fig. S2D). Despite being slightly different at these active promoters, H2Aub and H3K27me3 distributions in oocytes were largely overlapped at promoters. These results thus revealed similar landscapes of H2Aub and H3K27me3 in oocytes, in both broad distal domains and gene promoters.

### Maternal H2Aub but not H3K27me3 is erased from the broad distal domains after fertilization

To investigate how gamete H2Aub reprograms after fertilization, we further profiled H2Aub in zygotes as well as 2-cell, 4-cell, 8-cell, and blastocyst embryos by crossing two distinct parental strains of mice, namely PWK/PhJ (male) and C57BL/6N (female). The two biological replicates of each stage were highly reproducible (Table S1), we thus pooled data from replicates in subsequent analyses. As a control, we also included in our analyses previously published parental H3K27me3 data of gametes and early embryos (Zheng et al. 2016). In line with the observation that H2Aub is pervasively deposited in oocytes but not in sperms (Fig. 1A, 2A), zygotes showed a maternal-biased H2Aub enrichment (maternal/paternal reads ratio = 2.75) as revealed by our allelic analysis (Fig. S3A). However, such maternal bias was not observed in 2-cell, 4-cell, 8-cell, and blastocyst embryos (Fig. S3A), indicating a rapid reprogramming of gamete H2Aub towards parental equalization after fertilization. Indeed, H2Aub signals at H3K27me3 marked PMDs appeared to be inherited from oocytes by zygotes and then erased in embryos of later stages (Fig. 2A, S3B). In contrast, oocyte inherited H3K27me3 signals persisted in these PMDs throughout the preimplantation development (Fig. 2A, S3B). We further compared parental H2Aub and H3K27me3 signals across all H3K27me3 marked PMDs by plotting their average signals. Within the maternal genome, while similar enrichment was found for both H2Aub and H3K27me3 in zygotes, only H3K27me3 was enriched in the later-stage embryos, where H2Aub instead showed no apparent enrichment (Fig. 2B). Consistently, examination of maternal/paternal signal ratio in these PMDs showed that maternal-biased distribution was present only in zygotes for H2Aub but persisted in early embryos for H3K27me3 (Fig. 2C). Thus, our data demonstrate that H2Aub is decoupled from H3K27me3 in oocyte PMDs during preimplantation development.

**Figure 2.**
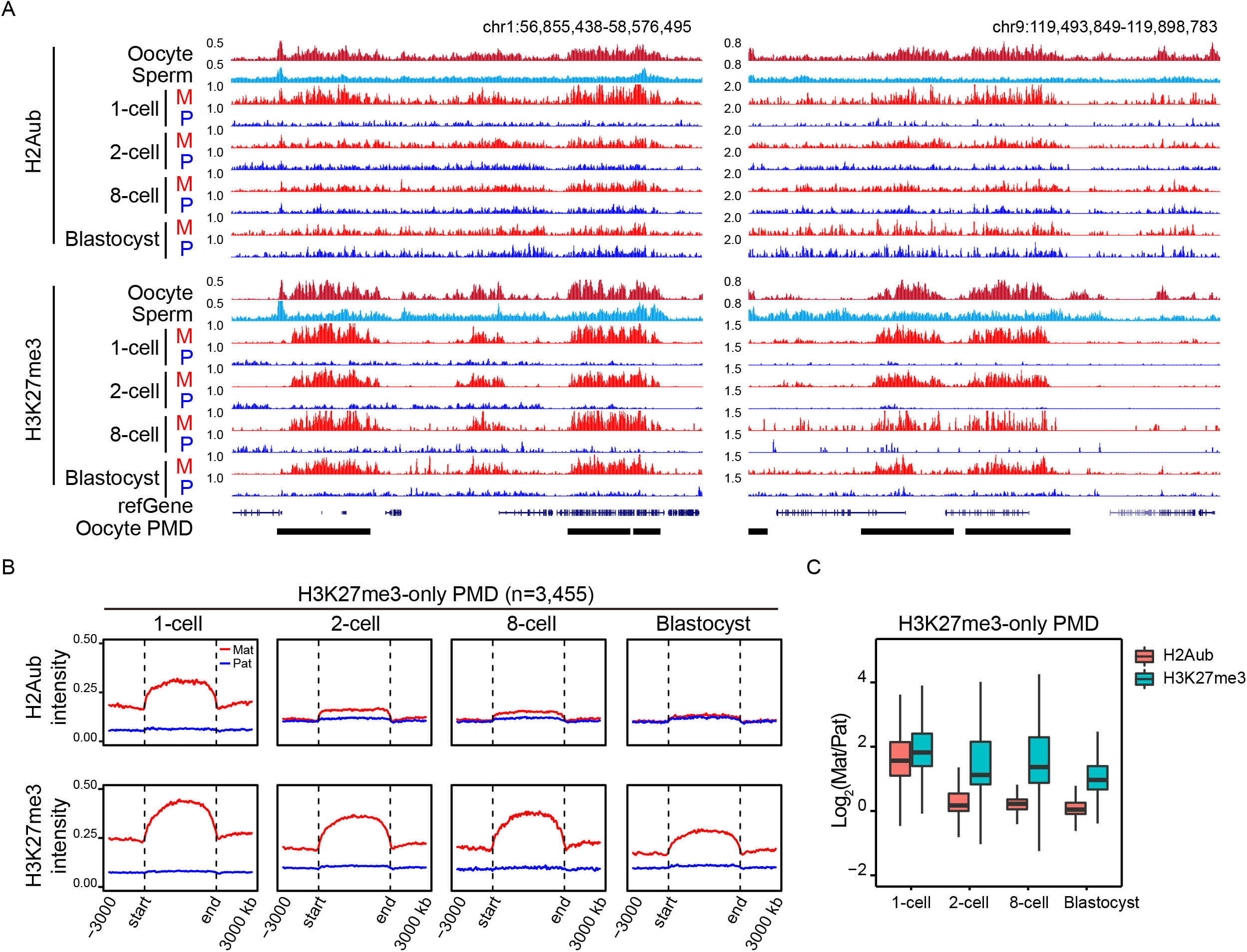
Maternal H2Aub but not H3K27me3 is erased from the broad distal domains after fertilization. **(A)** Genome browser view showing H2Aub and H3K27me3 enrichment in gametes and parental alleles of early embryos near oocyte PMDs. **(B)** Average H2Aub and H3K27me3 signals across H3K27me3-only PMDs in parental alleles of early embryos. **(C)** Box plot showing the maternal/paternal signal ratio of H2Aub and H3K27me3 in H3K27me3-only PMDs.

### H2Aub is not required for the maintenance of non-canonical imprinting

Maternal-biased deposition of H3K27me3 mediates non-canonical imprinting during preimplantation development (Inoue et al. 2017), and we have previously identified 4,135 large maternal-biased H3K27me3 domains in morula stage mouse embryos (Matoba et al. 2018). To investigate whether H2Aub and H3K27me3 function together in the maternal repression of non-canonical imprinting genes, we compared maternal and paternal signals in these maternal-biased H3K27me3 domains. As expected, H3K27me3 exhibited remarkably maternal-biased deposition in these domains at all preimplantation stages we examined (Fig. 3A, 3B). However, maternal bias of H2Aub in these domains was only observed in zygotes but not at later stages including 2-cell, 8-cell, and blastocyst embryos (Fig. 3A, 3B). Further examination of the 76 reported non-canonical imprinting genes (Inoue et al. 2017) also revealed similar decoupling of maternal H2Aub and H3K27me3 beyond the zygote stage (Fig. 3C, 3D, S3C). The absence of maternal H2Aub deposition across non-canonical imprinting genes thus demonstrated an irrelevance of H2AK119ub1 to the maternal repression of non-canonical imprinting genes during preimplantation development.

**Figure 3.**
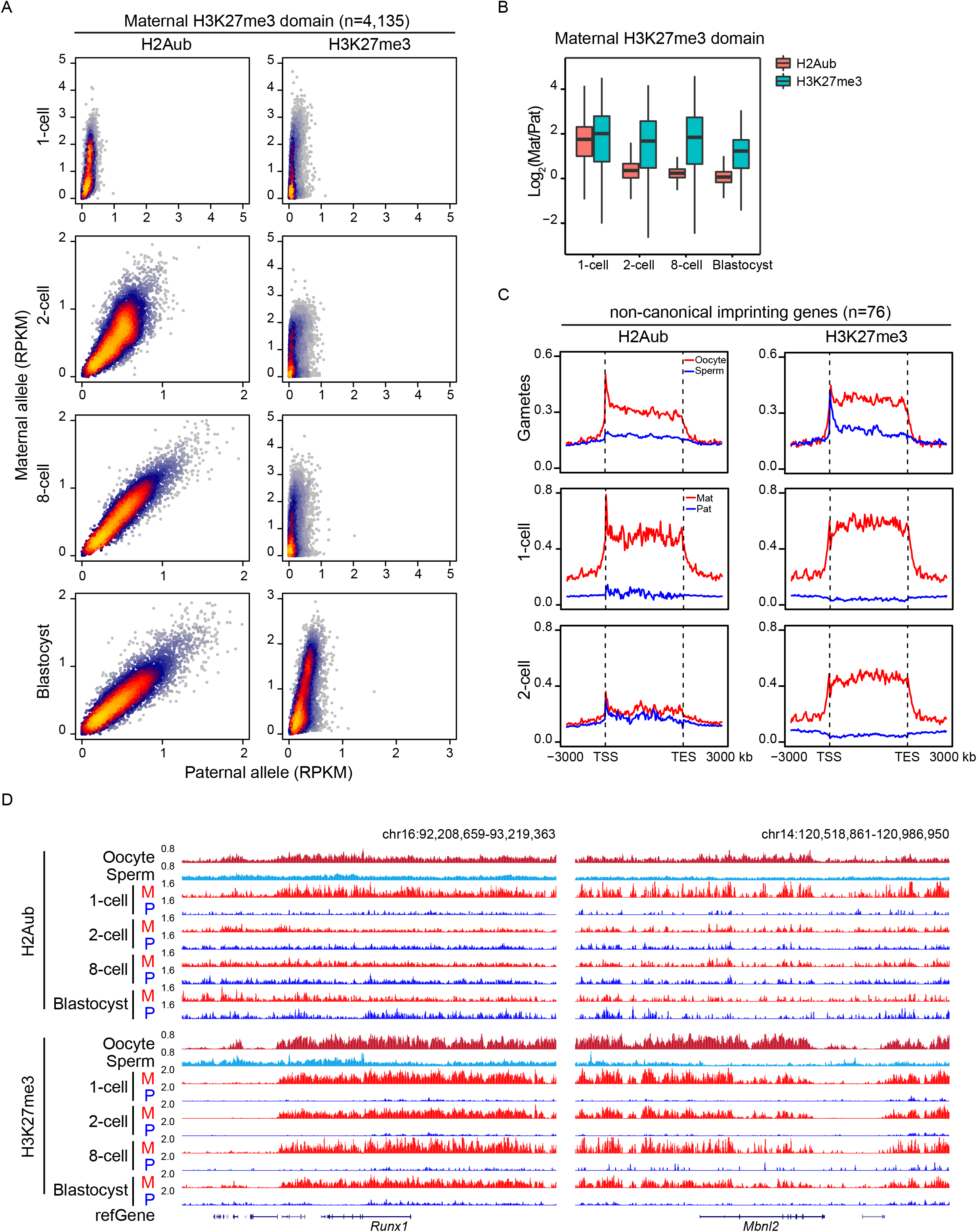
H2Aub is not required for the maintenance of non-canonical imprinting. **(A)** Density scatter plots comparing paternal and maternal ChIP-Seq signals in maternal H3K27me3 domains. **(B)** Box plot showing the maternal/paternal signal ratio of H2Aub and H3K27me3 in maternal H3K27me3 domains. **(C)** Average H2Aub and H3K27me3 signals across non-canonical imprinting genes in gametes and parental alleles of early embryos. **(D)** Genome browser view showing H2Aub and H3K27me3 enrichment in gametes and parental alleles of early embryos near *Runx1* (left) and *Mbnl2* (right).

### H2Aub is enriched at PcG-target promoters in the absence of H3K27me3 during early development

The decoupling of H2Aub and H3K27me3 in maternal H3K27me3 domains in early embryos promoted us to explore whether canonical PcG targets may still have similar deposition of H2Aub and H3K27me3 in early embryos as in mESCs. Consistent with the absence of bivalency during preimplantation development (Zheng et al. 2016; Xu and Xie 2018), H3K27me3 was actually not enriched at promoters of PcG target genes in 1-cell, 2-cell, and 8-cell embryos (Fig. 4A, 4B, S4A–S4C). But unexpected, we observed a persisting enrichment of H2Aub at these promoters during early development (Fig. 4A, 4B, S4A–S4C). Indeed, allelic analysis of Cluster 2 promoters, at which both H2Aub and H3K27me3 were enriched in oocytes, showed that H2Aub, but not H3K27me3, persisted in the maternal genome after fertilization (Fig. 4C), suggesting that H2Aub alone might be responsible for silencing these future bivalent genes during preimplantation development when H3K27me3 is not present. Consistent with this hypothesis, depletion of Ring1/Rnf2 (the catalytic components of PRC1), but not Eed (a core component of PRC2), resulted in a significant derepression of Cluster 2 genes in oocytes (Fig. 4D). Interestingly, instead of being inherited from sperms, paternal H2Aub enrichment at these promoters was gradually established towards the maternal levels after fertilization (Fig. 4C and S4A). This unexpected maintenance of H2Aub at PcG targets in early embryos was in stark contrast to H3K27me3, which are enriched in gametes but depleted from PcG targets upon fertilization (Fig. 4A–4C, S4A–S4C). Therefore, these data further revealed an early-embryo-specific decoupling of H2Aub and H3K27me3 at canonical PcG targets, where strong overlaps between H2Aub and H3K27me3 have been observed in mESCs and other cell types.

**Figure 4.**
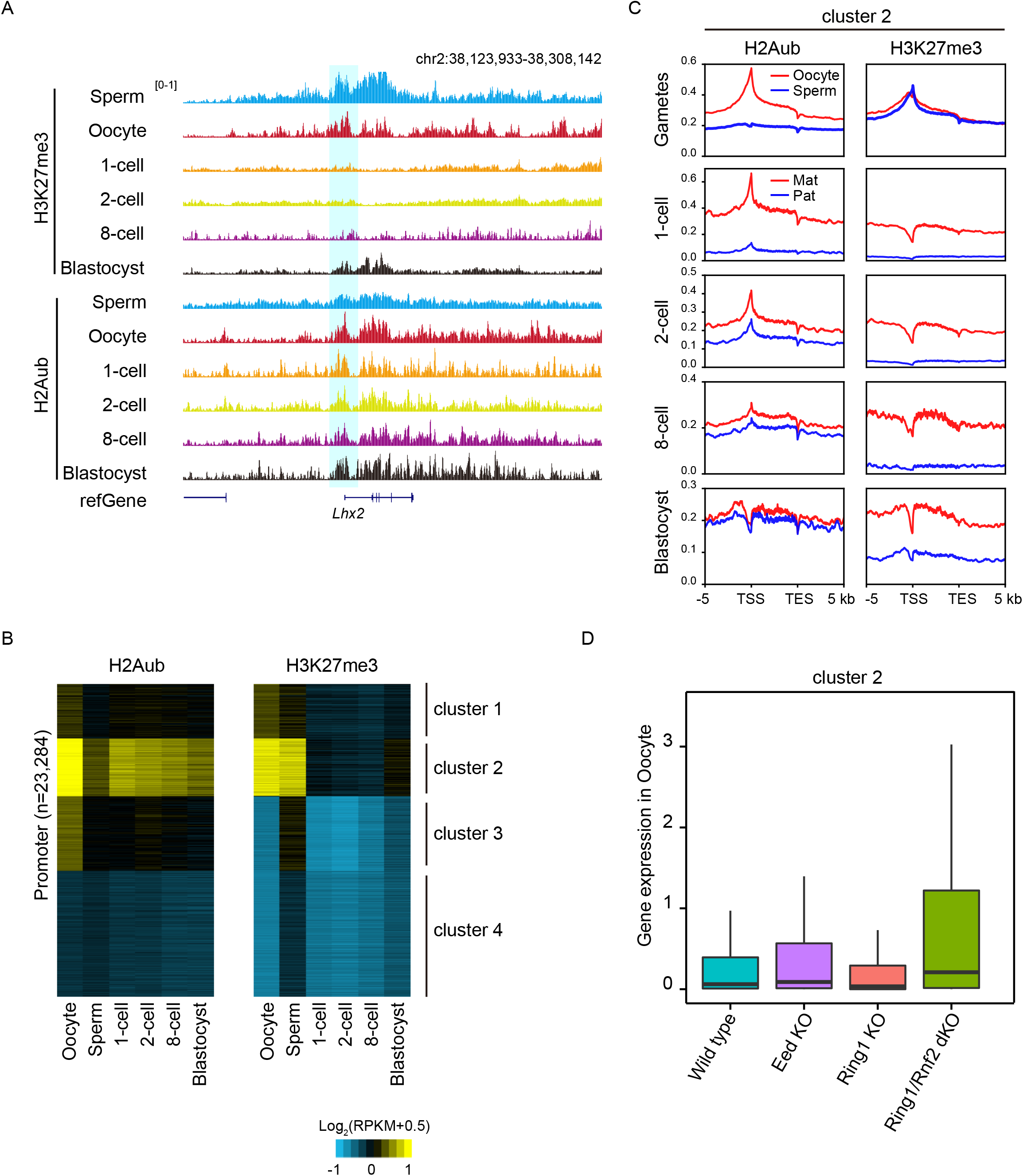
H2Aub is enriched at PcG-target promoters in the absence of H3K27me3 during early development. **(A)** Genome browser view showing H2Aub and H3K27me3 enrichment in gametes and early embryos near *Lhx2*. **(B)** Heatmaps showing the enrichment of H2Aub and H3K27me3 at promoter regions in gametes and early embryos. Clusters are the same as those in Figure 1D. **(C)** Average H2Aub and H3K27me3 signals across Cluster 2 genes in gametes and parental alleles of early embryos. **(D)** Box plot showing gene expression levels of Cluster 2 genes in wild-type, Eed-KO, Ring1-KO, and Ring1/Rnf2-dKO oocytes. KO, knockout; dKO, double knockout.

Collectively, our results show that H2Aub and H3K27me3, and thus PRC1 and PRC2, are unexpectedly decoupled during preimplantation development, with H2Aub enriched at canonical PcG targets and H3K27me3 placed in non-canonical-imprinting-related maternal domains. H2Aub appears to be responsible for repressing the expression of future bivalent genes in the absence of H3K27me3 in early embryos, prior to bivalency establishment. H3K27me3 alone is required for silencing the maternal allele of non-canonical imprinted genes. Our findings also raise important questions regarding the complex regulatory crosstalk between PRC1 and PRC2. For example, how the two complexes cooperate during oogenesis to deposit both H2Aub and H3K27me3 at canonical PcG targets and in broad distal domains respectively? Given that both H2Aub and H3K27me3 are present at these two types of regions in oocytes, PRC1 and PRC2 appear to function together at the establishment stage, by either hierarchical or mutual recruitment. Then how do they switch to the solo mode and function separately after fertilization? Do specific deubiquitinases or demethylases contribute to the decoupling of H2Aub and H3K27me3? While further investigations are required to answer these questions, it is important to note that such a genome-wide decoupling of the two closely connected histone marks in early embryos has never been observed in other cells. Thus, our findings not only provide an explicit example showing H2Aub and H3K27me3 can play distinct and independent roles, but also establish mouse early embryo as a unique model to study the independent recruitment and functions of PRC1 and PRC2 and their crosstalk.

## Materials and Methods

### Sample collection

PWK/PhJ and C57BL/6N mice were maintained under SPF conditions in a controlled environment. Experimental procedures and animal care were in accordance with the Animal Research Committee guidelines of Zhejiang University. Sperm were isolated from the caudal epididymis of male mice (10-12 weeks old) using a swim-up procedure. To collect oocytes, female mice (21-23 days old) were injected with 5 IU of pregnant mare serum gonadotropin (PMSG) and humanely euthanized 44 h later. Fully grown oocytes were harvested from ovaries in M2 medium (Sigma-Aldrich). To collect early embryos, C57BL/6N female mice (21-23 days old) were injected with 5 IU of PMSG followed by human chorionic hormone (hCG) 44 h later. Superovulated female mice were mated with 10-12-week-old PWK/PhJ males. Successful mating was confirmed by the presence of vaginal plugs. Zygotes were harvested from oviducts at 28 h after hCG injection and then cultured in KSOM Mouse Embryo Media (Millipore) at 37°C with 5% CO2, and 2-cell, 4-cell, 8-cell, and blastocyst stage embryos were harvested at 16 h, 30 h, 48 h, and 72 h after culture, respectively. Only embryos showing the correct developmental stage were collected. Five blastocysts, and 200-400 oocytes or 1-cell to 8-cell embryos were used for CATCH-Seq.

### CATCH-Seq library preparation and sequencing

CATCH-Seq was performed essentially as ULI-NChIP-Seq (Brind’Amour et al. 2015) with the addition of synthetic double-stranded carrier DNA prior to immunoprecipitation and in wash buffers. The carrier DNA was prepared by annealing the forward strand (/5’NH2C6G/TAGGGATAACAGGGTAATTAGGGATAACAGGGTAATTAGGGATAACAGGGTAATTAGGGATAACAGGGTAATTAGGGATAACAGGGTAATTAGGGATAACAGGGTAAT*/3’ddC/) and the reverse strand (/5’NH2C6G/ATTACCCTGTTATCCCTAATTACCCTGTTATCCCTAATTACCCTGTT ATCCCTAATTACCCTGTTATCCCTAATTACCCTGTTATCCCTAATTACCCTGTTAT CCCTA*/3’ddC/). Each strand has an amino modifier with a C6 spacer arm at the 5’ end and a dideoxyribose nucleotide at the 3’ end to block adaptor ligation, and asterisks represent phosphorothioate bonds. To completely remove small amount of carrier DNA that might be ligated to adaptors due to potential synthesis errors, pre-amplified libraries were digested by I-SceI before final amplification. Briefly, cells were pelleted and resuspended in nuclear isolation buffer (10 mM Tris-HCl, pH8.0, 140 mM NaCl, 5 mM MgCl2, 0.6% NP-40, 0.1% Triton X-100, 0.1% deoxycholate, 1 x EDTA-free proteinase inhibitor cocktail and 1 mM PMSF). Chromatin was then fragmented with MNase (2 U/μl at 21 °C for 7.5 min for cultured cells; 2 U/μl at 37 °C for 7.5 min for fully grown oocytes and early embryos; 8 U/μl at 25°C for 10 min for sperms) and diluted in NChIP immunoprecipitation buffer (20 mM Tris-HCl pH 8.0, 2 mM EDTA, 150 mM NaCl, 0.1% Triton X-100, 1 × EDTA-free protease inhibitor cocktail and 0.1 mM PMSF). Carrier DNA (30 ng) was added into the fragmented chromatin to reduce DNA loss during immunoprecipitation. The fragmented chromatin was then pre-cleared with 5 μl of Dynabeads Protein A/Protein G (ThermoFisher) and immunoprecipitated overnight at 4 °C with the pre-prepared antibody-beads complex, which contains 700 ng H3K4me3 antibody (15410003, Digagenode) or H2Aub antibody (8240, Cell Signaling Technology) and 5 μl Dynabeads Protein A/Protein G. After immunoprecipitation, beads were washed twice with 150 μl of low salt wash buffer (20 mM Tris-HCl, pH 8.0, 0.1% SDS, 1% Triton X-100, 2 mM EDTA, 150 mM NaCl, 1 x EDTA-free proteinase inhibitor cocktail, 0.1 mM PMSF, and 0.2 ng/μl carrier DNA) and twice with 150 μl of high salt wash buffer (20 mM Tris-HCl, pH 8.0, 0.1% SDS, 1% Triton X-100, 2 mM EDTA, 500 mM NaCl, 1 x EDTA-free proteinase inhibitor cocktail, 0.1 mM PMSF, and 0.2 ng/μl carrier DNA). Protein-DNA complexes were eluted from beads in 100 μl ChIP elution buffer (100 mM NaHCO3 and 1% SDS) for 2 h at 65 °C. Immunoprecipitated DNA was purified by phenol-chloroform extraction followed by ethanol precipitation. Adaptor ligation was performed using the NEBNext Ultra II DNA Library Prep Kit for Illumina (E7645, New England Biolabs), and library pre-amplification was performed by 9 cycles of PCR. The pre-amplified library was digested by I-SceI (5 U/μl; 2 h incubation at 37 °C followed by 20 min heat inactivation at 65 °C). Barcoded final libraries were generated by further amplification of the carrier-DNA-removed pre-amplification products and were sequenced on the Illumina HiSeq X Ten platform.

### ChIP-Seq data analysis

H3K4me3/H3K27me3 ChIP-Seq data in mouse gametes/embryos and H2Aub data of mESC were obtained from published data sets (Zhang et al. 2016; Zheng et al. 2016; Li et al. 2017). CBX7 and RYBP ChIP-seq data in mESCs were obtained from published data sets (Healy et al. 2019). For both published data sets and the data sets generated in this study, reads were trimmed using Trim Galore (v0.4.4) with default parameters and aligned against the mouse genome build mm9 using Bowtie2 (v2.3.4.1) with default parameters. All unmapped reads and PCR duplicates were removed. Peaks were analyzed using MACS2 (v2.1.1.20160309) with the parameters “-q 0.1 --broad-cutoff 0.1 --nomodel --nolambda --broad --extsize 300 -B --SPMR -g mm”. Signal tracks for each sample were generated using the “wigToBigWig” utility from UCSC and visualized using UCSC genome browser. Correlation coefficient was generated using deeptools (v2.5.4).

### Allele assignment of sequencing reads

Uniquely mapped and PCR duplicates removed reads were used to assign to parental origins. SNP information between PWK/PhJ and C57BL/6N mouse strains was obtained from the Mouse Genomes Project. Reads covering SNP sites were extracted for allele assignment. When multiple SNPs were present in a read, the parental origin was determined by votes from all SNPs and the read was assigned to the allele that had at least two thirds of the total votes. For downstream analysis, RPKM (reads per kilobase of bin, per million mapped reads) values were computed for each allele, by counting the numbers of allelic reads per kilobase of bin per million of SNP trackable reads.

### Clustering and heatmap analysis

Reads in promoters regions (TSS+/-2 kb), oocyte PMDs, and maternal H3K27me3 domains were calculated using the “coverage” command in bedtools (v2.26.0) and then normalized to RPKM. The k-means clustering of H2AK119ub1 and H3K27me3 enrichment at promoters in oocytes was conducted using “kmeans” function in R. To generate heatmaps showing ChIP-Seq signal enrichment around TSS or PMDs, TSS+/- 5kb regions or 2x PMD regions were divided into 40 bins and enrichment level in each bin was computed as RPKM and further normalized using Z-score normalization. Average intensity profiles were generated using in-house R scripts. Density scatter plots were generated using “LSD” package in R. Heatmaps were shown using Java Treeview. Genes marked by H3K27me3 at their promoters in mESCs were defined as PcG target genes (Zheng et al. 2016).

### RNA-seq data analysis

RNA-Seq data of wild-type, Eed-KO, Ring1-KO, and Ring1/Rnf2-DKO oocytes were obtained from published data sets (Zhang et al. 2016; Du et al. 2020). Raw reads were trimmed to 50 bp and mapped to the mouse genome (mm9) using TopHat (v2.1.1) with default parameters. Only uniquely mapped reads were kept for downstream analysis. The RNA abundance of each gene was quantified using Cufflinks (v2.2.1).

### Data availability

ChIP-Seq data sets generated in this study have been deposited in the Gene Expression Omnibus (GEO) database under the accession number GSE169199.

## Author contributions

L.S. conceived and supervised the project; Y.Z. performed data analysis and generated figures; J.Y. developed and performed CATCH-Seq; Y.R. and Y.W. collected the samples; L.S. and Y.Z. wrote the manuscript with inputs from H.F., and all authors discussed the results and commented on the manuscript.

## Acknowledgments

We are grateful to all members in the Fan laboratory and the Shen laboratory for helpful discussion. This project was supported by National Natural Science Foundation of China (32022023, 31871478), National Key Research and Development Programs of China (2017YFC1001500), and Zhejiang Provincial Natural Science Foundation of China (LR18C060001).

## Supplementary Figure Legends

**Figure S1.**
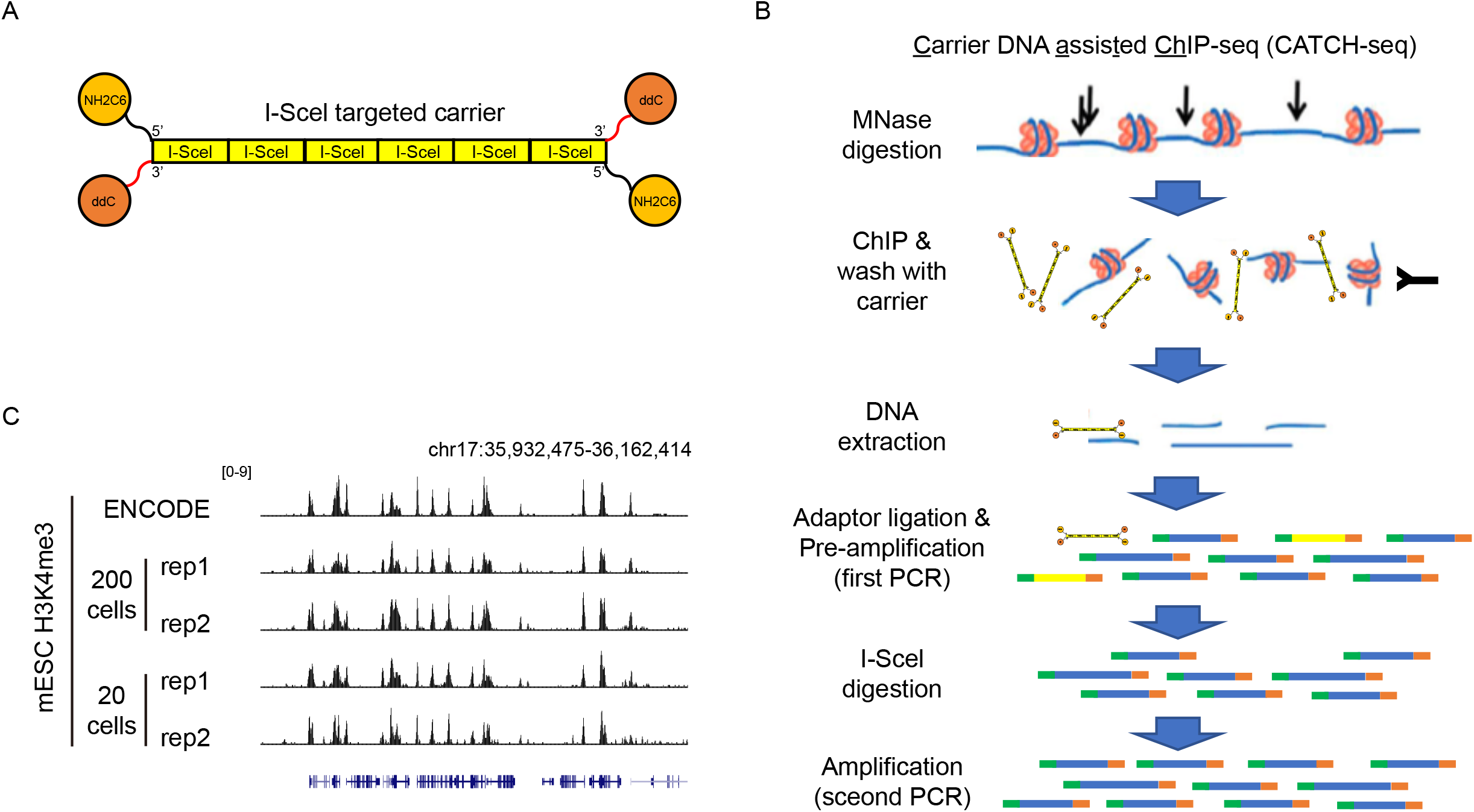
Validation of CATCH-Seq in mESCs. **(A)** A schematic of the carrier DNA used in CATCH-Seq. NH2C6 is a 5’ amino modifier with a C6 spacer arm; ddC is a dideoxyribose nucleotide; red lines at the 3’ end, phosphorothioate bonds; I-SceI, the 18-bp recognition site of the meganuclease I-SceI. **(B)** Schematic illustration of CATCH-Seq. The carrier DNA generally cannot be ligated to adaptors because of the 5’ amino modifier and 3’ dideoxyribose nucleotide, and small amount of carrier DNA being ligated to adaptors due to potential synthesis error will be removed by I-SceI digestion after library pre-amplification. **(C)** Genome browser view comparing mESC H3K4me3 signals generated by CATCH-Seq using indicated numbers of input cells, and the signal generated by bulk ChIP-Seq (data downloaded from ENCODE).

**Figure S2.**
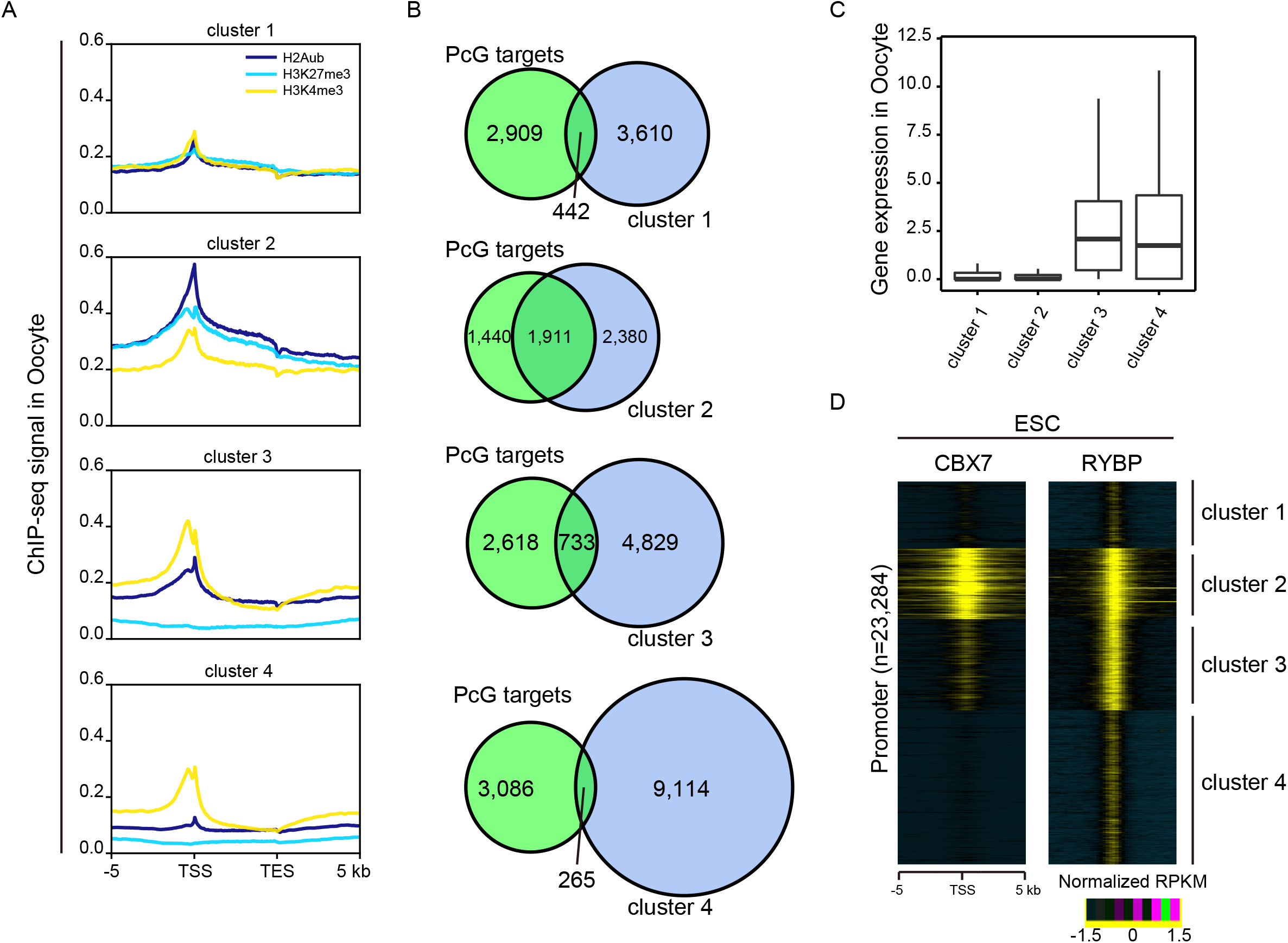
H2Aub and H3K27me3 enrichment at gene promoters in oocytes. **(A)** Average H2Aub, H3K27me3, and H3K4me3 signals across four clusters of genes in oocytes. Genes are clustered into four groups using k-means clustering based on promoter H2Aub and H3K27me3 signals in oocytes, as in Figure 1D. **(B)** Venn diagrams showing overlap of PcG target genes with the four clusters of genes. **(C)** Box plot showing expression levels of the four clusters of genes in oocytes. **(D)** Heatmaps showing the enrichment of CBX7 and RYBP around TSS of the four clusters of genes in mESC.

**Figure S3.**
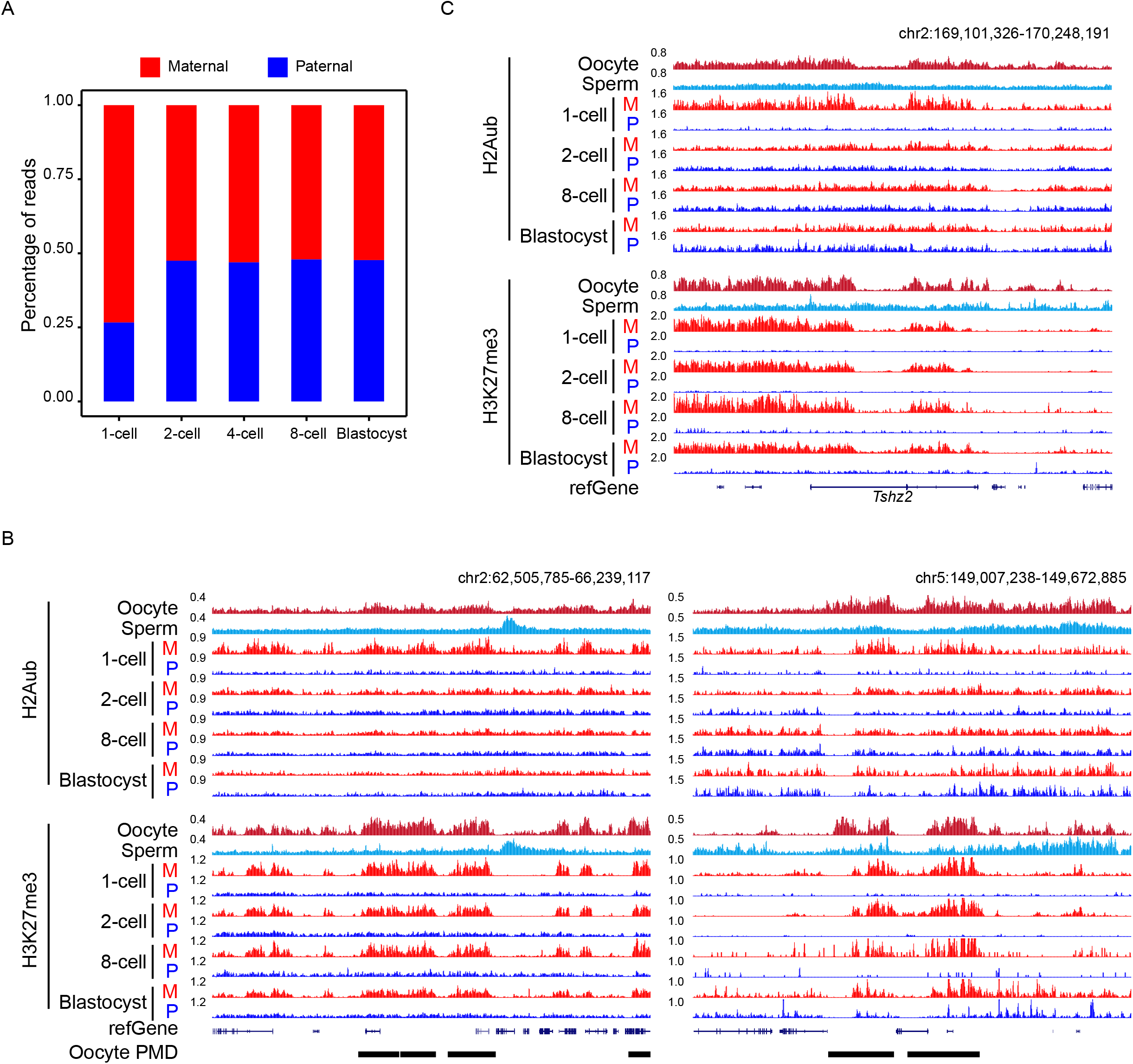
Allelic analysis of H2Aub deposition in maternal H3K27me3 domains. **(A)** Bar plot showing percentage of parental H2Aub ChIP-seq reads in early embryos. **(B)** Genome browser view showing H2Aub and H3K27me3 enrichment in gametes and parental alleles of early embryos near oocyte PMDs. **(C)** Genome browser view showing H2Aub and H3K27me3 enrichment in gametes and parental alleles of early embryos near *Tshz2*.

**Figure S4.**
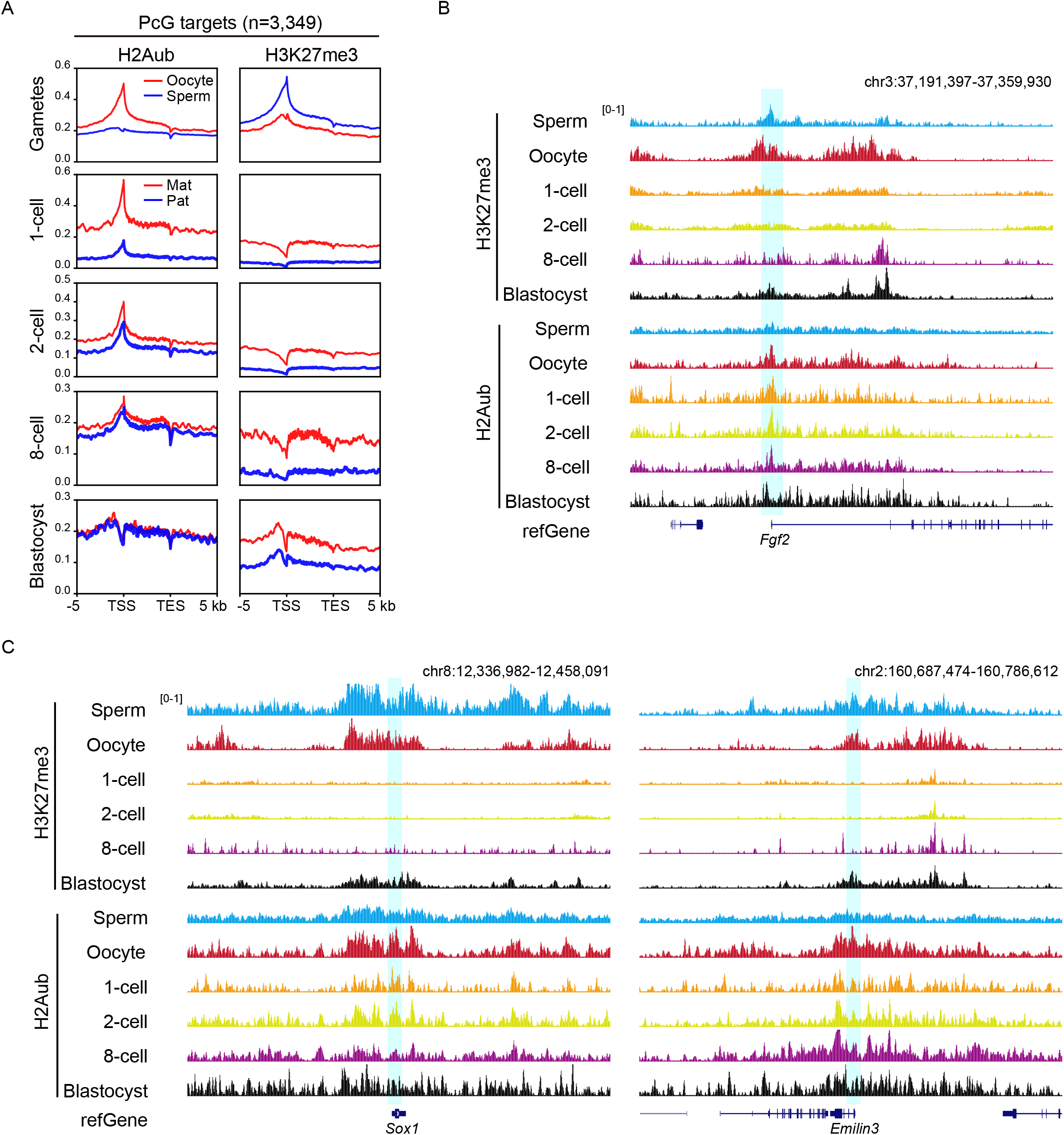
Allelic analysis of H2Aub and H3K27me3 at PcG targets. **(A)** Average H2Aub and H3K27me3 signals across PcG target genes in gametes and parental alleles of early embryos. **(B)** Genome browser view showing H2Aub and H3K27me3 enrichment in gametes and embryos near *Fgf2*. **(C)** Genome browser view showing H2Aub and H3K27me3 enrichment in gametes and embryos near *Sox1* (left) and *Emilin3* (right).

## Supplementary Tables

**Table S1.** Summary of datasets generated in this study.

## Notes

### Competing Interest Statement

The authors have declared no competing interest.

